# Robust EEG brain-behavior associations emerge only at large sample sizes

**DOI:** 10.64898/2026.02.06.704323

**Authors:** Nicolas Le Guern, Sahar Allouch, David Pascucci, Aida Ebadi, Mahmoud Hassan

## Abstract

Electroencephalography (EEG) offers unique access to the oscillatory dynamics that shape human cognition and behavior. Yet reported associations between EEG features and behavioral measures often fail to replicate, raising fundamental concerns about the robustness of this literature. A key reason is the predominance of small sample sizes (median ∼30 participants), but the consequences for reproducibility have not been systematically evaluated. Here, we provide the first large-scale empirical test of this key issue using the largest available harmonised resting-state EEG dataset, comprising 2,292 participants across healthy individuals and clinical populations. We extracted a comprehensive set of spectral, temporal, complexity, and dynamical features, and assessed their correlations with behavioral, cognitive, and mental health measures. Using extensive resampling across a wide range of sample sizes, we show that small samples produce unstable associations with inflated and inconsistent effect sizes, whereas larger cohorts converge on smaller but reproducible effects. These findings establish sample size as a critical determinant of reliability in EEG research and call for a re-evaluation of how oscillatory dynamics are linked to human cognition. By moving beyond low-powered correlations toward large-scale, reproducible approaches, the field will be better positioned to uncover the true role of neural oscillations in shaping behavior.

## Introduction

Neural oscillations, first described almost a century ago, are now recognized as a fundamental organizing principle of brain function. Recorded non-invasively with electroencephalography (EEG), they provide a powerful window into the temporal coordination of neural activity that underlies cognition and behavior. Oscillatory dynamics have been implicated across a wide range of domains, including perception^1–3^, attention^4–7^, memory^8–11^, decision-making^12–15^, motor control^16– 19^, sleep^20–22^, vigilance^23–25^, and emotion^26–28^, and are increasingly studied in relation to behavioral performance, mental states, and clinical outcomes^29–35^. A central framework in this work was the study of brain-behavior associations, in which EEG features are linked to cognitive or clinical measures as a means of probing the functional relevance of neural rhythms.

Despite their appeal, most reported EEG brain-behavior associations, including those proposed as candidate biomarkers, have proven difficult to replicate^36–39^. Similar concerns have been documented in other domains of neuroscience. In MRI, large-scale brain-wide association studies (BWAS) have shown that small samples can produce inflated and unreliable effects, sparking a broader debate about the standards required for robust brain research^40^. These challenges may be even more acute in EEG, a modality characterized by high sensitivity to noise, variability across sites, and a literature dominated by small-sample studies (median ∼30 participants). Yet, although the impact of sample size on brain-behavior associations has been systematically examined in MRI, no comparable large-scale evaluation has been carried out for EEG oscillatory features, despite their central role in cognitive and clinical neuroscience.

Here, we provide the first systematic large-scale evaluation of sample size effects on EEG brain-behavior associations. We leverage a harmonized, multisite resting-state dataset comprising 2,292 participants, spanning both healthy and clinical populations. From these recordings, we extracted a comprehensive set of EEG features spanning spectral, temporal, complexity, and dynamical domains. Using repeated subsampling (n = 1,000), we systematically varied sample sizes from as few as 10 participants to several hundred, and quantified how this influenced both the statistical significance of EEG-behavior correlations and the stability of effect size estimates. Our analyses raise critical questions about the robustness of many findings currently reported in the field, situate EEG within the broader reproducibility debate, and provide empirical guidance for designing more rigorous and sustainable approaches to studying oscillatory mechanisms in cognition and clinical neuroscience.

## Results

### Correlations at small sample size are unstable in direction, magnitude, and significance

We examined how sample size affects the reliability of EEG brain-behavior associations by analyzing resting-state EEG data from 2,292 participants aggregated across four harmonized datasets (see Methods). To this end, we implemented systematic resampling across a wide range of sample sizes, starting from as few as 10 participants. For each target sample size, we drew subsamples 1,000 times without replacement and computed Spearman correlations between EEG-derived features and behavioral measures spanning demographics, cognition, and mental health. This procedure was repeated across all sample sizes, with correlations estimated both for individual scalp channels and for channel-averaged features.

At smaller sample sizes, EEG-behavior associations were highly variable, with both the sign and the strength of correlations fluctuating across iterations. As an illustrative example, we show the relationship between Beta Relative Power at channel F4 and a behavioral measure of Depression (CBCL Withdrawn). At *n* = 30 (reflecting the typical sample size of many EEG studies), the observed correlation alternated between positive, negative, and near-zero across the 1,000 resamples. This pattern contrasts sharply with the full dataset, which revealed a small near zero correlation (Fig. 1a). Similar instability was seen at the spatial level: the set of scalp channels yielding significant correlations changed substantially across resamples and often bore little resemblance to the topographic distribution observed in the full sample (Fig. 1b). These patterns generalized across all feature-behavior associations: correlations at *n* = 30 differed markedly from those observed in the full sample in both magnitude and direction, consistently across the entire feature space (Fig. 1c).

**Fig. 1.**
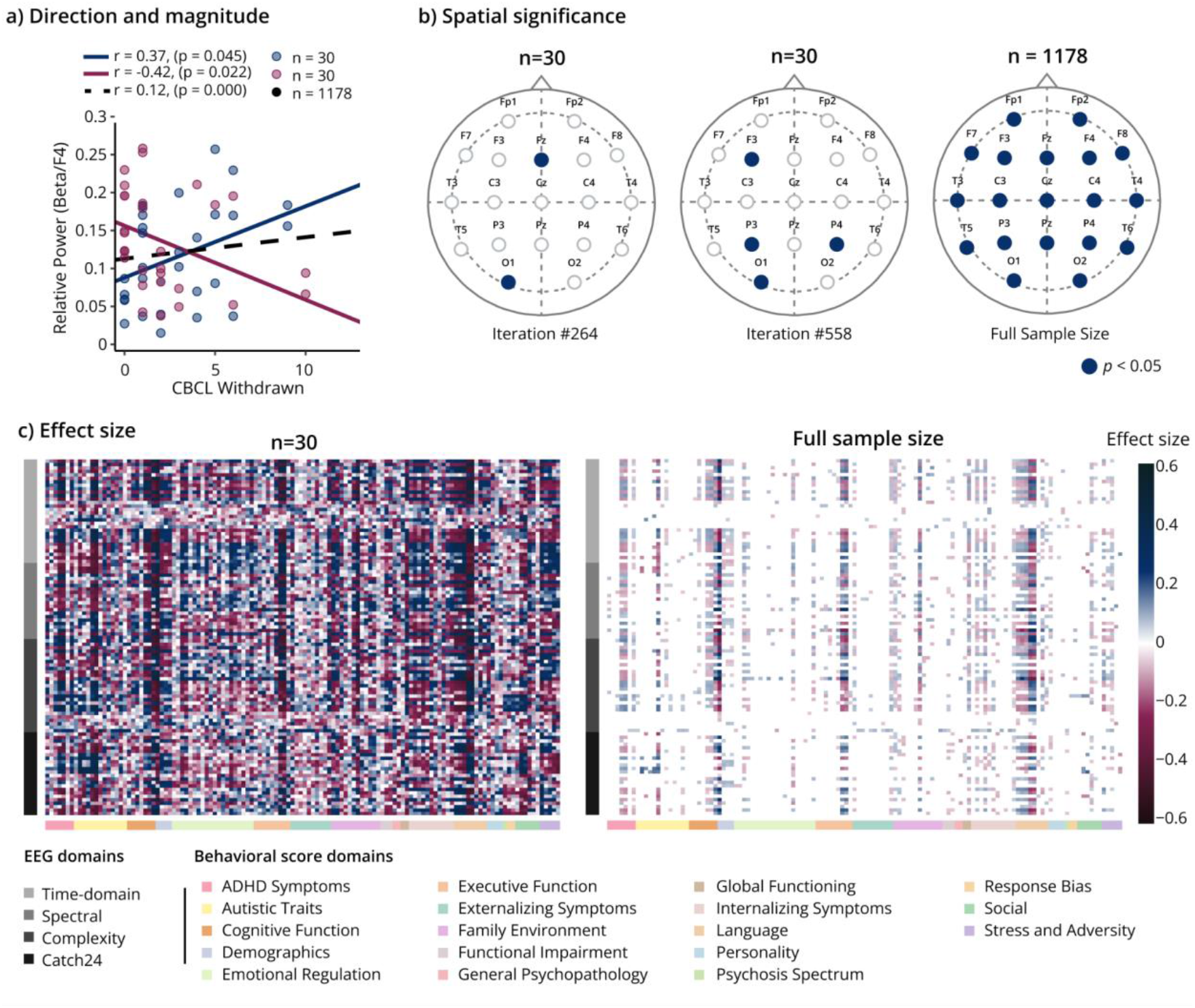
Instability of EEG-behavior correlations at small sample size (n = 30). **a**, Two independent random subsets of 30 participants from the ADHD group yield strong but opposing correlations, in contrast to the near-zero association observed in the full dataset (black dashed line). **b**, Spatial distribution of significant correlations in two example subsampling iterations (out of 1000), showing little resemblance to the full-sample topography. **c**, Heatmaps of correlation coefficients (r) between all 103 EEG features (averaged over all channels) and behavioral scores. At *n*=30, values reflect the mean of significant correlations across 1,000 iterations, shown alongside the full-sample pattern.

### Consistency emerges at larger sample sizes

We sought to characterize the consistency with which each association reached statistical significance across increasing sample sizes. For each EEG-behavior pair, we calculated a significance percentage, defined as the proportion of 1,000 iterations in which the correlation yielded *p* < 0.05. The sample size threshold was then defined as the minimum sample size at which this percentage exceeded 95%. Across all associations, we identified four distinct reproducibility profiles (Fig. 2a-d). For associations significant at the full sample size, the significance percentage increased monotonically with *n* and eventually crossed the 95% threshold (Fig. 2a). Among associations that were not significant in the full dataset, three distinct patterns were observed: a gradual increase that never reached the 95% threshold (Fig. 2b), a consistently low significance percentage across sample sizes (Fig. 2c), and a progressive decline that stabilized near zero (Fig. 2d).

**Fig. 2.**
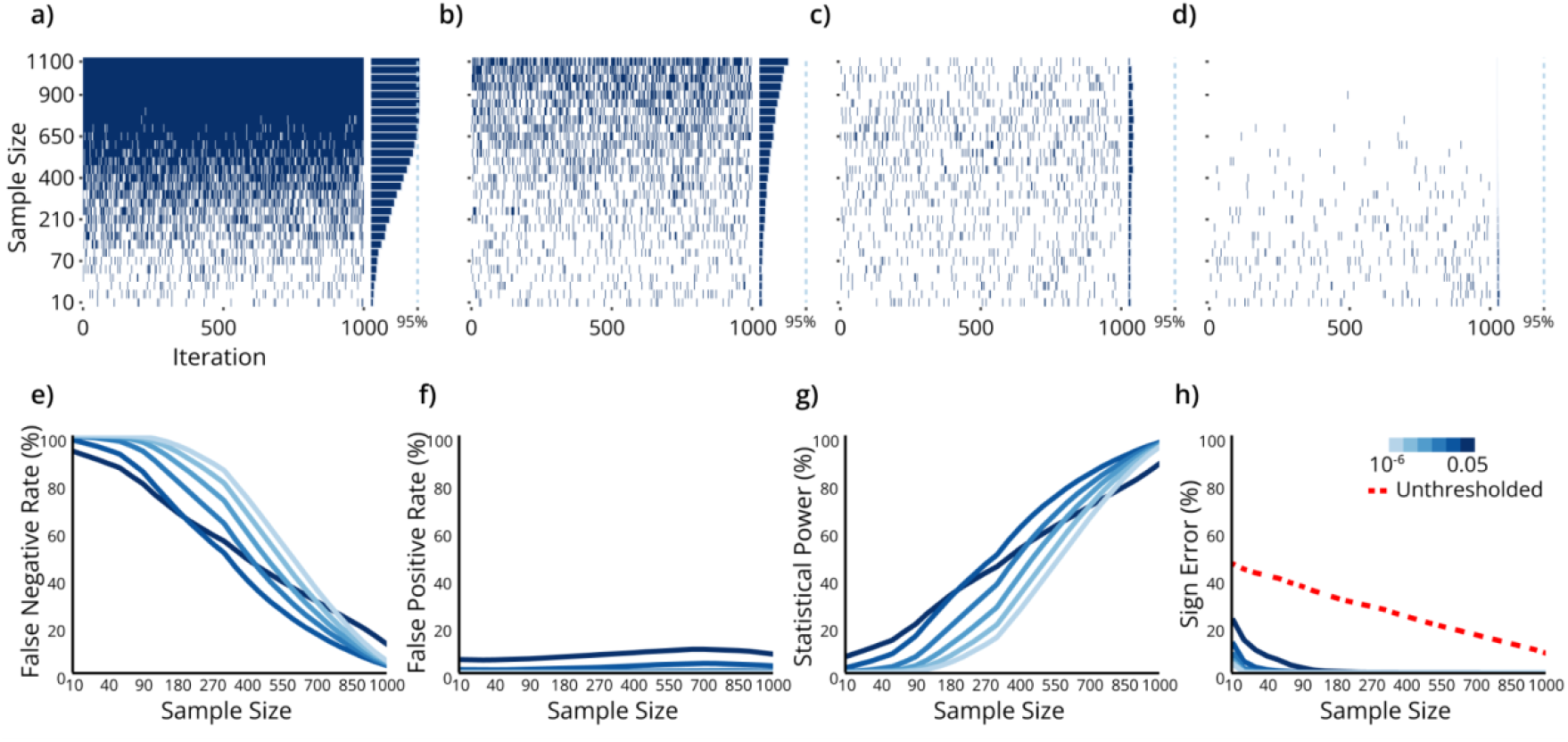
Patterns of significance and error properties of correlations across sample sizes for ADHD. **a–d**, Heatmaps illustrating representative patterns of feature-score associations across sample sizes and iterations. Each row corresponds to a given sample size, and each column to one subsampling iteration (dark blue = significant, white= non-significant). Four reproducibility profiles were observed: **a**, associations that steadily reached the 95% reproducibility threshold; **b**, associations with increasing but subthreshold significance; **c**, unstable associations with irregular significance across sample sizes; and **d**, associations with consistently low significance. **e–h**, Global error metrics as a function of sample size. **e**, False negative rate (%), **f**, False positive rate (%), **g**, Statistical power, **h**, Sign error rate (%). The red dashed line indicates the sign error rate without applying a significance threshold. Results in **e-h** are based on feature-score pairs with at least 1000 subjects available.

For significant associations, reproducibility thresholds were substantially higher than the sample sizes typically used in EEG correlation studies: 202 ± 69 for HC, 808 ± 290 for ADHD, and 288 ± 85 for ASD. Considerable spatial variability was also evident: even within a single EEG feature– score association, different scalp channels often required markedly different sample sizes to reach reproducibility (Fig. S2,3).

When examining global error metrics (including false negatives, false positives, statistical power, and sign errors), we observed a strong dependence on sample size (Fig. 2e–h). False negatives predominated at small sample sizes, reaching nearly 90% under uncorrected thresholds and close to 100% after multiple-comparison correction. These rates declined steadily with increasing *n*, dropping to 11.8% (uncorrected) and 3.91% (corrected) at *n*=1,000. False positives showed only a modest increase and never exceeded 10%, while sign errors were largely restricted to very small subsamples (*n*<150). Applying a more stringent significance threshold further increased false negatives while decreasing both false positives and sign errors. Overall, this profile of rapidly increasing sensitivity, modest false-positive inflation, and sharply reduced false negatives was consistent across diagnostic groups. Although Fig. 2e–h illustrates results for ADHD, the same patterns were observed in HC and ASD. (see Supplementary Fig. S4).

### Effect sizes at lower sample sizes are exaggerated

Beyond statistical significance, the magnitude of EEG–behavior associations is critical for evaluating their scientific and clinical relevance. Small sample sizes yielded highly unstable correlations, with both strength and sign varying markedly between iterations (Fig. 3b). For instance, at *n* = 30, significant associations frequently exhibited correlations exceeding ±0.4, with the smallest detected effect size at 0.36, yet these effects diminished substantially at larger sample sizes (Fig. 3c,d). Across all significant associations in ADHD, approximately 80% of correlations at *n* = 30 were inflated by at least 200% relative to estimates derived from the full dataset (Fig. 3a), with similarly high levels of inflation observed for ASD and HC (Fig. S4).

**Fig. 3.**
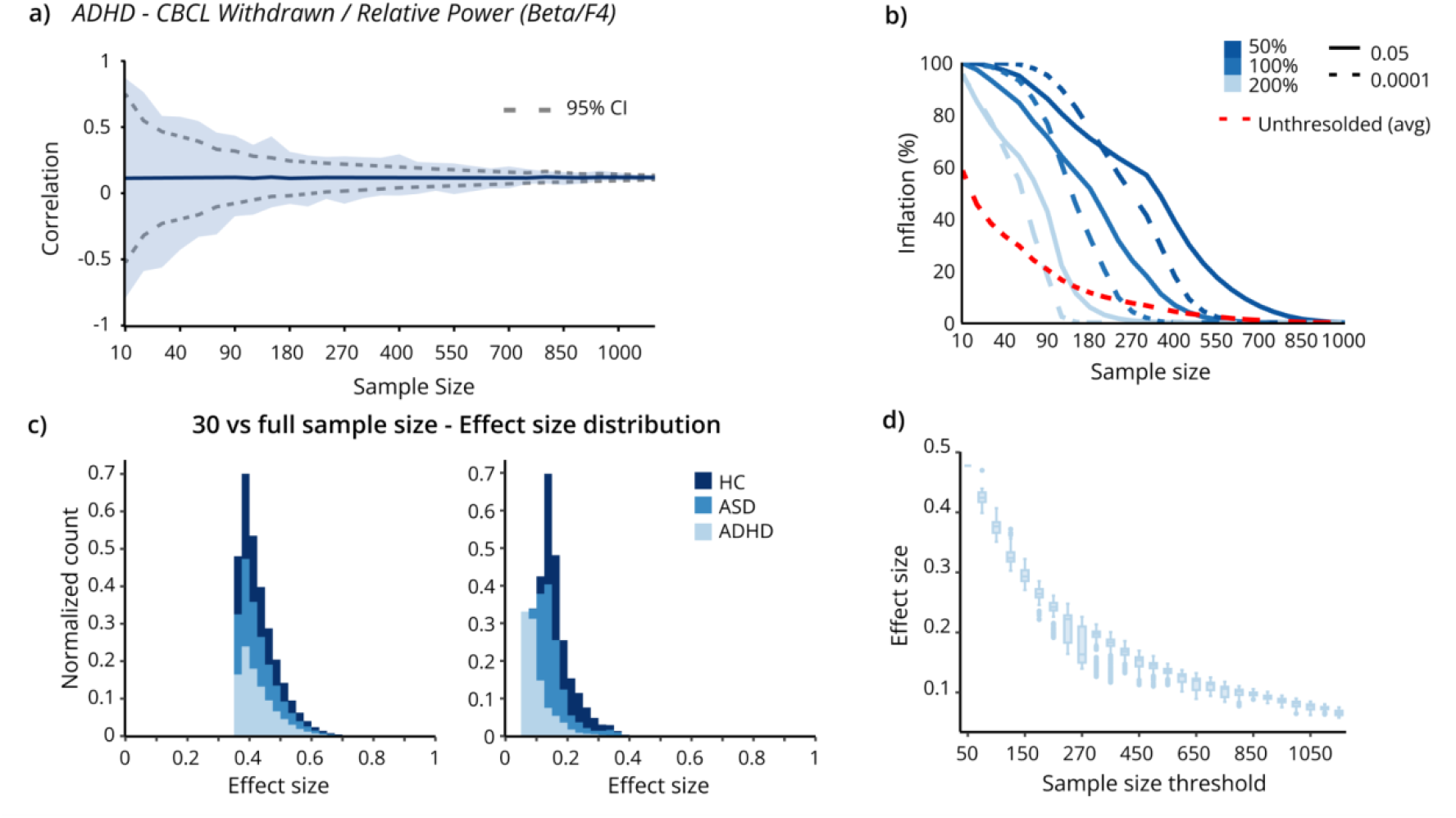
Impact of sample size on effect size estimates. **a**, Correlation estimates (±95% CI) for the association between CBCL Withdrawn and relative Beta power at channel F4 across subsamples. **b**, Proportion of inflated effect sizes in ADHD across sample sizes, shown under different significance thresholds (p < 0.05, p < 0.0001) and inflation levels (50%, 100%, 200%). The red dashed line denotes the average effect size inflation without applying a significance threshold. **c**, Distribution of average effect sizes at n = 30 (left) compared to the full sample (right). **d**, Effect sizes at the full sample size plotted against sample size thresholds for the ADHD group.

This inflation is a direct consequence of limited statistical power: when sample sizes are small, only the strongest (and often spurious) correlations cross the significance threshold, biasing the set of observed effects toward overestimates. As *n* increases, the range of significant correlations narrows, and the variability across iterations declines. At the maximum available sample size, correlations converged to small values, rarely exceeding |0.25| (ADHD: 1.29%; ASD: 5.15%; HC: 11.34% of the significant associations, Fig. 3c). Notably, this value is lower than the minimum effect sizes observed at *n*=30. This pattern was evident in both healthy and clinical groups, underscoring that inflated effect sizes are a general property of low-*n* EEG-behavior studies rather than a feature of specific populations.

### Adjusting for age attenuates brain-behavior correlations

Initial analyses revealed that the strongest associations between brain measures and behavioral scores were predominantly driven by age, accounting for 3.40% of all significant correlations. Recognizing that age and sex can act as confounding factors in EEG-behavior relationships, we repeated the analyses using partial correlations, controlling for age and sex. This adjustment substantially reduced the proportion of significant associations (11.83% vs. 3.20%) and yielded consistently smaller correlation coefficients. In ADHD, effect sizes no longer exceeded |0.25|, while in ASD and HC, the proportion of significant associations with |*r*| > 0.25 decreased to 0.75% and 6.77%, respectively (Fig. 4a,c, and supplementary Fig. S6). The median sample size threshold of partial correlations increased for ADHD from 850 to 1,000 compared with unadjusted correlations, while the overall distribution retained a similar shape. The main difference was a reduction in the number of significant features. Patterns of statistical significance, inference metrics, and inflation remained qualitatively consistent. Notably, at *n* = 30, false negative rates were higher (93%), and inflation reached 97%, while the overall pattern was preserved (see Supplementary Fig. S5,6). The relationship between sample size thresholds and effect sizes exhibited the same trend, but overall effect sizes were smaller, indicating that many apparent associations were driven by age.

**Fig. 4.**
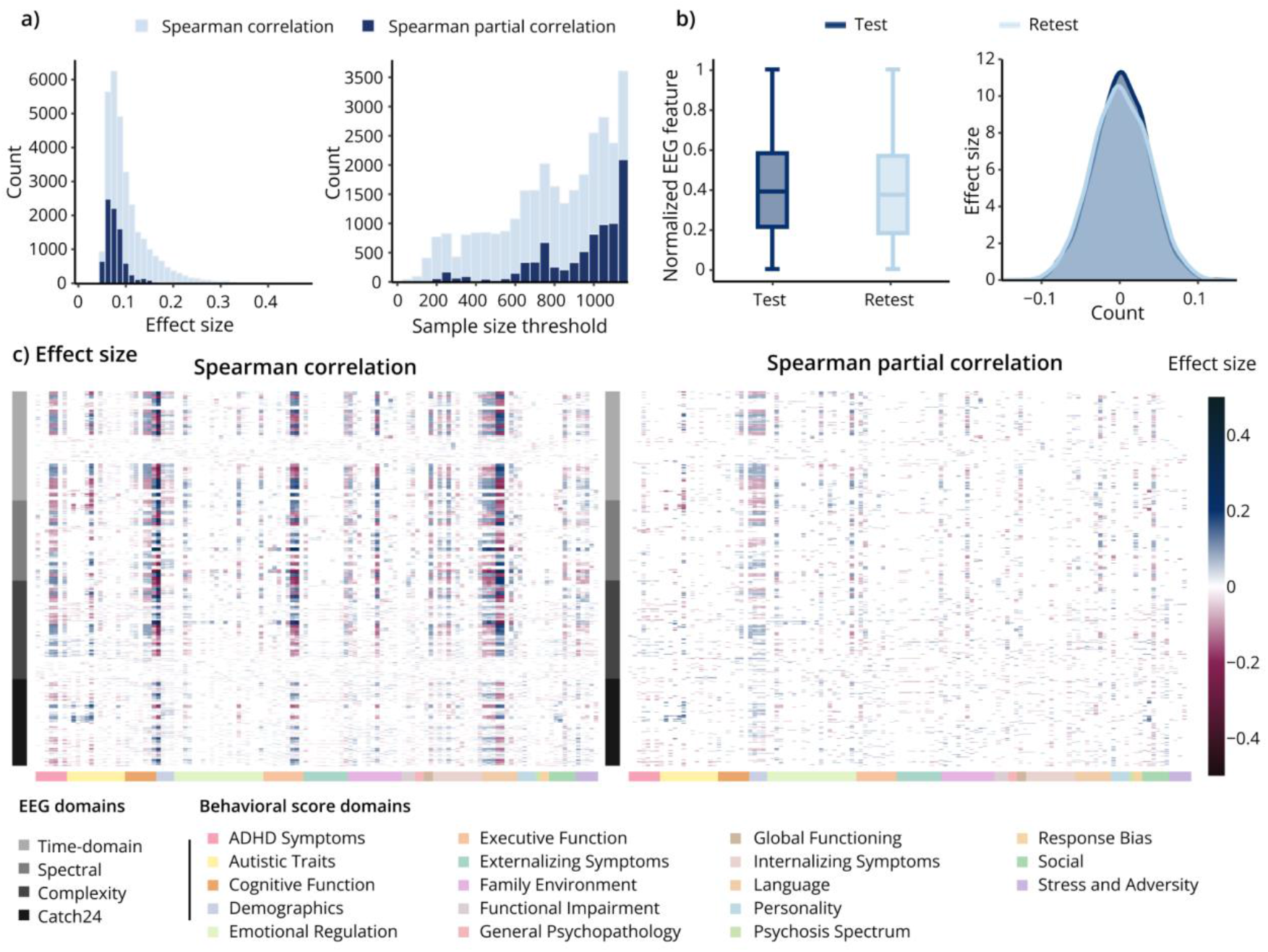
Control analyses: effects of covariates and intra-subject variability on associations. **a**, Distribution of effect sizes at full sample (left) and sample size thresholds (right) for Spearman correlation and Spearman partial correlation. **b**, Box plot of normalized EEG features (left) and distribution of effect sizes (right) derived from two independent, non-overlapping epoch sets. **c**, Heatmaps of significant effect sizes for Spearman correlations (left) and Spearman partial (right) correlations across all associations.

### Analysis outcomes remain stable across independent epochs

To evaluate whether the choice of epochs influenced our feature estimates and downstream analyses, we computed two independent feature sets from the same EEG recordings by randomly selecting five non-overlapping epochs per set. Analyses were restricted to channel-averaged features within the ADHD group. Feature distributions were highly consistent across the two sets (test: median = 0.39; retest: median = 0.37), and the effect-size distributions of feature–score correlations were nearly identical (Fig. 4b). Importantly, the sample size thresholds required for significance remained stable (test: median = 1000; retest: median = 1000), indicating that epoch selection does not materially affect the results.

## Discussion

Oscillatory brain activity, as measured by EEG, has long been considered a fundamental substrate of brain-behavior associations, offering mechanistic insights into how neural dynamics shape brain function and dysfunction. Yet reported associations have been strikingly inconsistent across studies^36–39^. By systematically resampling a large multi-site EEG dataset with 1,000 iterations across varying sample sizes, we show that the stability of observed brain-behavior associations is critically constrained by sample size. This type of analysis has become feasible only with the recent availability of large-scale, harmonized EEG datasets, which provide both the statistical power to assess stability across a wide range of sample sizes and the diversity across sites and participants needed to generalize reproducibility estimates beyond single cohort^41^. Our findings parallel concerns raised in other neuroimaging modalities, particularly MRI-based brain-wide association studies, where small samples have been shown to yield inflated and unreliable effects^40^. Together, this convergence indicates that the reproducibility challenges observed here are not unique to EEG but reflect a broader issue across cognitive and clinical neuroscience.

At sample sizes typical of the EEG literature (∼30 participants), the correlations fluctuated markedly across iterations and failed to approximate the estimates derived from the full cohort. In some cases, this instability persisted across all sample sizes tested; in others, increasing sample size led to more consistent outcomes. Significant associations observed in the full dataset generally required substantially larger subsamples -few hundreds of participants-to achieve stability, far exceeding the typical sample sizes used in EEG research. At smaller sample sizes, false negatives were frequent, reflecting a high likelihood of missing genuine effects, and sign errors were common and only declined once sample sizes exceeded roughly 150 participants. These results underscore that small samples not only obscure true effects but also undermine replicability. Rather than prescribing a minimum sample size for reliable inference, our study emphasizes the magnitude of sampling error and the inferential challenges that arise in commonly studied cohorts. Even samples of 500 participants may still inadequately represent the broader population, particularly in light of well-documented recruitment biases^42^.

The distribution of effect sizes revealed a systematic bias: smaller samples tended to inflate estimated magnitudes, whereas larger samples more accurately reflected the subtle associations present in the data. As sample size increased, even weak effects reached statistical significance, but without the inflation observed in smaller samples. This pattern illustrates the dual pitfalls of insufficient sample size: i) diminished sensitivity to true effects and ii) overestimation of those that are detected. Importantly, this bias arises from a well-known selection mechanism: in small-*n* settings, only those features with spuriously large estimated effects are likely to surpass the significance threshold, leading to a biased subset of reported associations^40^. This “winner’s curse” illustrates how reliance on Type-I error control as the primary evidentiary standard perpetuates a literature skewed toward overstated findings and inconsistent replication when sample sizes are inadequate. Beyond sample size alone, covariates such as age and sex substantially shaped the observed associations. When comparing Spearman correlations with partial correlations that controlled for these covariates, we found fewer and weaker significant effects at the full sample level, with the strongest Spearman correlations largely driven by age. This indicates that developmental variation (5-18 years) exerts a dominant influence on EEG features, underscoring the necessity of accounting for confounds alongside adequate sample size.

Several methodological aspects should be considered when interpreting these findings. First, although our study draws on one of the largest resting-state EEG datasets available, the diagnostic groups were limited to ADHD and ASD. This choice reflects the availability of sufficiently large-scale EEG data (>500 participants) with accompanying behavioral measures, but may restrict the scope of our analyses. The statistical principle that small samples yield unstable associations is likely to generalize across populations. However, the extent of instability and the sample sizes required to achieve reproducible estimates may differ in other clinical conditions (such as schizophrenia, depression, mood disorders, or neurodegenerative disorders), which can exhibit distinct effect size distributions, sources of variability, and levels of heterogeneity. Second, while our iterations-based approach systematically quantifies variability across sample sizes, it cannot completely eliminate potential biases related to participant recruitment, diagnostic criteria, or site-specific differences. Third, as sample size approaches the maximum available in a group, subsampling variability decreases because participants are repeatedly included across iterations. Although this does not alter the core conclusions of our study, namely, the instability of small-sample findings, it may influence precise estimates of reproducibility thresholds.

Beyond these methodological considerations, it is important to reflect on broader questions concerning the analytical frameworks used to study brain-behavior associations. A limitation of the present work is its reliance on a univariate correlation framework. While this approach reflects common practice in EEG research and allows for a transparent evaluation of reproducibility, it does not capture the complexity of multivariate relationships. The univariate framework implicitly assumes one-to-one mappings between neural features and behavioral measures, whereas psychological phenomena are more plausibly supported by many-to-one mappings, with multiple neural configurations giving rise to the same outcome^43^. Consequently, a single EEG feature may overlook genuine associations that operate through alternative neural pathways and may lack specificity if the same signal reflects different functions across contexts. Moreover, in psychiatry and neuroscience it is common to find studies where a significant correlation between an EEG feature and a clinical score is presented as evidence of a biomarker. Yet a group-level association does not imply that the same feature can predict outcomes at the level of an individual^44^. For a measure to qualify as a biomarker, it must demonstrate robustness, reproducibility, and predictive validity, as well as clinical or functional relevance (for example, the capacity to track symptom changes) beyond what can already be inferred from inexpensive behavioral assessments^41^.

Importantly, this study should not be taken as a call to abandon small-n designs. Looking ahead, progress in EEG-behavior research is likely to follow two complementary paths. Large-scale consortia will be essential for detecting small but reproducible cross-sectional associations at the population level, with findings that may primarily inform public health and social policies^44^. Small-n studies, by contrast, excel at maximizing signal-to-noise and isolating large effects that would be diluted in broad, heterogeneous samples. They can focus on patient groups with pronounced brain-behavior differences, employ within-subject or longitudinal frameworks to trace dynamics over time, and use controlled experimental manipulations to test mechanisms directly^44^. Such designs provide insights into individual trajectories of change and intervention responses, questions that large-scale studies are poorly suited to address.

In summary, our results reveal the profound impact of sample size on the reproducibility of EEG brain-behavior associations. Small samples produce unstable and inflated effects, while large cohorts converge on small but reliable correlations. By demonstrating this empirically in a large multi-site dataset, we highlight a fundamental reproducibility challenge in EEG research that must be addressed for the field to advance.

## Methods

### Dataset

The EEG data used in this study were drawn from the same source as our previous studies^45,46^, comprising 2292 subjects (age: 9.96 ± 3.01, 39.75% male) with healthy controls (n=529) and subjects diagnosed with ASD (n=542) and ADHD (n=1221). Data were aggregated from four datasets: the Healthy Brain Network Dataset (*[HBN](https://fcon_1000.projects.nitrc.org/indi/cmi_healthy_brain_network/index.htm)*)^47,48^, Multimodal Resource for Studying Information Processing in the Developing Brain (*MIPDB*)^47^, Autism Biomarker Consortium for Clinical Trials Dataset (*ABCCT*)^49^, Multimodal Developmental Neurogenetics of Females with ASD (*femaleASD*)^50^.

### Behavioral scores

The behavioral measures available across the four datasets spanned a broad range of domains. Altogether, 53 scores were distributed across 18 domains, yielding 254 distinct subscores. To minimize statistical bias from underpowered analyses and to ensure robust and generalizable estimates, subscores with fewer than 200 participants were excluded. This resulted in a final set of 152 subscores across 18 categories (Table S1). Because not all scores were present in every dataset, the number of participants contributing to each subscore varied both within and between groups (Fig. S1). Ultimately, we retained 127 subscores for ADHD, 44 for ASD, and 77 for HC.

### EEG preprocessing

Resting-state EEG was originally recorded with high-density 128-channel systems under eyes-open conditions and subsequently downsampled to 19 channels using the standard 10–20 montage. As source localization was not performed in this study, high spatial resolution was deemed unnecessary, and channel reduction allowed for substantial computational efficiency. The EEG preprocessing pipeline adhered to the automated approach described in our recent studies^45,46^. Subjects with channel interpolation rates exceeding 20% were excluded after preprocessing, after which data underwent visual inspection to remove excessively noisy recordings. Finally, a subject-wise z-transformation was applied: for each subject, the mean across all channels was subtracted and the result divided by the across-channel standard deviation, thereby normalizing EEG amplitudes, ensuring comparability.

### EEG features

We extracted 103 EEG features reflecting a broad spectrum of signal characteristics (Table S2 provides the full list of computed features). These included conventional frequency-domain measures such as absolute and relative power-computed across canonical frequency bands (delta [δ], 1–4 Hz; theta [θ], 4-8 Hz; alpha [α], 8-13 Hz; beta [β], 13-30 Hz; gamma [γ], 30-45 Hz), as well as features capturing temporal dynamics and signal complexity. Notably, we included metrics from the CAnonical Time-series CHaracteristics (Catch22) toolbox, a curated set of 22 features selected from an initial pool of 4,791 for their discriminative power across a wide range of classification tasks, along with the mean and standard deviation of the signal, resulting in 24 time-series features in total^51^. By incorporating both widely used and less conventional descriptors, our feature set was designed to capture the multifaceted nature of EEG signals beyond standard spectral analyses.

Given the multi-site nature of our dataset, we applied feature harmonization using neurCombat^52^ to reduce site-related variability while preserving biologically relevant signals. In our implementation, we specified age as a continuous covariate and included sex and group as categorical covariates to ensure that their contributions to feature variability were preserved during harmonization.

### Correlation framework: repeated subsampling analysis

A subsampling procedure was implemented to assess the stability of correlation estimates across varying sample sizes. Subsample sizes increased in variable steps depending on group size, with finer resolution at small *n* and larger steps at higher *n* (see Supplementary Table S3). For each subsample size, *n* subjects were randomly drawn without replacement from the full cohort, and associations between EEG features and behavioral scores were computed using Spearman’s rank correlation^53^ implemented in Python via the *SciPy* library^54^. This procedure was repeated 1,000 times per sample size. The sample size threshold was defined as the smallest *n* at which correlations reached statistical significance in more than 95% of iterations. To reduce computational demands, the number of iterations was capped at 100 once this threshold was exceeded.

To account for confounding effects of age and sex, both known to influence EEG activity and cognitive development^55–58^, we also computed Spearman partial correlations. This approach isolates the unique contribution of each EEG feature to the behavioral or demographic measure of interest while removing shared variance attributable to age and sex.

### Inference error metrics

We computed four inference error metrics as a function of sample size: false negatives, false positives, statistical power, and sign errors^40^. “True significance” was defined according to correlations observed in the full dataset. For each subsample size, false negatives were defined as the proportion of iterations in which a feature that was significant in the full dataset failed to reach significance in the subsample. Conversely, false positives were defined as the proportion of iterations in which a feature that was not significant in the full dataset was nevertheless identified as significant in the subsample. Statistical power was defined as one minus the false negative rate. Sign errors were defined as instances in which the direction of the association in a subsample differed from that in the full dataset, and were computed for both thresholded and unthresholded *p*-values. Figures in the main text depict averages of these metrics across all feature–score pairs. Significance thresholds (10-6 to 0.05) were adjusted for multiple comparisons using the Bonferroni method.

### Inflation

Effect size inflation was quantified for associations identified as significant by dividing the effect size obtained at each significant subsample *n* by the effect size estimated from the full sample. For each sample size, we calculated the proportion of cases in which this ratio exceeded 50%, 100%, or 200%, and these proportions were then averaged across all significant associations. This analysis was conducted at two p-value thresholds: *p* < 0.05 and *p* < 10^-4^.

### Intra-subject reliability

To assess the consistency and robustness of EEG–behavior associations, we evaluated intra-subject reliability. This analysis was performed in the ADHD group from the HBN dataset, which provided a sufficiently large sample. Continuous EEG recordings were segmented into non-overlapping 4-second epochs, from which two independent, non-overlapping sets of five epochs were randomly selected for each participant. Features were averaged within each set across epochs and then across channels to yield one value per EEG feature. Spearman’s rank correlations with cognitive scores were computed separately for the two sets using the same analysis pipeline as in the main study. To reduce computational demands, the number of subsampling iterations was limited to 500.

## Data availability

ABCCT and femaleASD datasets can be requested from the NIMH Data Archive platform (https://nda.nih.gov/edit_collection.html?id=2288, https://nda.nih.gov/edit_collection.html?id=2021); MIPDB and HBN datasets are accessible at Child Mind Institute website (https://fcon_1000.projects.nitrc.org/indi/cmi_eeg/index.html, https://fcon_1000.projects.nitrc.org/indi/cmi_healthy_brain_network/index.html).

## Code availability

Codes are available at https://github.com/MINDIG-1/EEG-CorrResampling-Framework. We used the *pingoiun*^59^ python package for statistical calculations, ‘*MNE*’ python package^60^ (https://mne.tools/stable/index.html) for EEG signal processing and custom python-based scripts for the remaining analysis and visualization.

## Acknowledgment

This work was funded by MINDIG as part of its R&D activity. It was also supported by the “Region Bretagne”, Inno R&D project no. 23001155 and Rennes Metropole (AICE project) and the INCR (PsyNorm and Creapark projects). We would like to thank all the researchers who shared their data in open-access and all the participants (patients and controls) who approved the use of their data in research.

## Authors contribution

A.E., N.L., M.H., and S.A. conceived the study and oversaw data analysis and interpretation. N.L. and A.E. drafted the manuscript and generated the figures, with critical revisions from all authors, including S.A., D.P., and M.H. S.A. assisted with data preprocessing and feature extraction. D.P. provided key feedback on the project and contributed specifically to the test-retest analysis.

## Ethics declaration

### Ethics approval

Ethical oversight was provided by the Chesapeake Institutional Review Board and Yale Institutional Board (Yale, SCRI). All procedures performed were in accordance with the ethical standards of the 1964 Helsinki Declaration and its later amendments or comparable ethical standards. All participants (or their legal guardians) provided written informed consent prior to inclusion in the study.

## References

1. Demiralp, T. et al. Gamma amplitudes are coupled to theta phase in human EEG during visual perception. Int. J. Psychophysiol. 64, 24–30 (2007).

2. Busch, N. A., Dubois, J. & VanRullen, R. The Phase of Ongoing EEG Oscillations Predicts Visual Perception. J. Neurosci. 29, 7869–7876 (2009).

3. Melloni, L. et al. Synchronization of Neural Activity across Cortical Areas Correlates with Conscious Perception. J. Neurosci. 27, 2858–2865 (2007).

4. Klimesch, W., Doppelmayr, M., Russegger, H., Pachinger, T. & Schwaiger, J. Induced alpha band power changes in the human EEG and attention. Neurosci. Lett. 244, 73–76 (1998).

5. Vandecappelle, S. et al. EEG-based detection of the locus of auditory attention with convolutional neural networks. eLife 10, e56481 (2021).

6. Li, G. et al. Drivers’ EEG Responses to Different Distraction Tasks. Automot. Innov. (2023) doi:10.1007/s42154-022-00206-z.

7. Deiber, M.-P. et al. Linking alpha oscillations, attention and inhibitory control in adult ADHD with EEG neurofeedback. NeuroImage Clin. 25, 102145 (2020).

8. Magosso, E. & Borra, D. The strength of anticipated distractors shapes EEG alpha and theta oscillations in a Working Memory task. NeuroImage 300, 120835 (2024).

9. Katerman, B. S., Li, Y., Pazdera, J. K., Keane, C. & Kahana, M. J. EEG biomarkers of free recall. NeuroImage 246, 118748 (2022).

10. McKeon, S. D. et al. Aperiodic EEG and 7T MRSI evidence for maturation of E/I balance supporting the development of working memory through adolescence. Dev. Cogn. Neurosci. 66, 101373 (2024).

11. Li, Y., Pazdera, J. K. & Kahana, M. J. EEG decoders track memory dynamics. Nat. Commun. 15, 2981 (2024).

12. Si, Y. et al. Predicting individual decision-making responses based on single-trial EEG. NeuroImage 206, 116333 (2020).

13. Golnar-Nik, P., Farashi, S. & Safari, M.-S. The application of EEG power for the prediction and interpretation of consumer decision-making: A neuromarketing study. Physiol. Behav. 207, 90–98 (2019).

14. Jacobs, J., Hwang, G., Curran, T. & Kahana, M. J. EEG oscillations and recognition memory: Theta correlates of memory retrieval and decision making. NeuroImage 32, 978–987 (2006).

15. Yau, Y. et al. Evidence and Urgency Related EEG Signals during Dynamic Decision-Making in Humans. J. Neurosci. 41, 5711–5722 (2021).

16. McFarland, D. J., Sarnacki, W. A. & Wolpaw, J. R. Electroencephalographic (EEG) control of three-dimensional movement. J. Neural Eng. 7, 036007 (2010).

17. Müller-Putz, G. R., Scherer, R., Pfurtscheller, G. & Rupp, R. EEG-based neuroprosthesis control: A step towards clinical practice. Neurosci. Lett. 382, 169–174 (2005).

18. Ang, K. K. et al. A Randomized Controlled Trial of EEG-Based Motor Imagery Brain-Computer Interface Robotic Rehabilitation for Stroke. Clin. EEG Neurosci. 46, 310–320 (2015).

19. Jerbi, K. et al. Inferring hand movement kinematics from MEG, EEG and intracranial EEG: From brain-machine interfaces to motor rehabilitation. IRBM 32, 8–18 (2011).

20. Campbell, I. G. EEG Recording and Analysis for Sleep Research. Curr. Protoc. Neurosci. 49, (2009).

21. Lambert, I. & Peter-Derex, L. Spotlight on Sleep Stage Classification Based on EEG. Nat. Sci. Sleep Volume 15, 479–490 (2023).

22. Gu, Y., Gagnon, J. & Kaminska, M. Sleep electroencephalography biomarkers of cognition in obstructive sleep apnea. J. Sleep Res. 32, e13831 (2023).

23. Babiloni, C., Güntekin, B., Yener, G. & Del Percio, C. qEEG Methods to Probe Abnormal Brain Rhythms Related to Quiet Vigilance in Patients with Dementia Due to Alzheimer’s, Parkinson’s, and Lewy Body Diseases. in Psychophysiology Methods (eds Valeriani, M. & De Tommaso, M.) vol. 206 67–89 (Springer US, New York, NY, 2024).

24. Olbrich, S. et al. EEG-vigilance and BOLD effect during simultaneous EEG/fMRI measurement. NeuroImage 45, 319–332 (2009).

25. Olbrich, S. et al. EEG Vigilance Regulation Patterns and Their Discriminative Power to Separate Patients with Major Depression from Healthy Controls. Neuropsychobiology 65, 188–194 (2012).

26. Li, X. et al. EEG Based Emotion Recognition: A Tutorial and Review. ACM Comput. Surv. 55, 1–57 (2023).

27. Gong, L., Li, M., Zhang, T. & Chen, W. EEG emotion recognition using attention-based convolutional transformer neural network. Biomed. Signal Process. Control 84, 104835 (2023).

28. Li, Y. et al. GMSS: Graph-Based Multi-Task Self-Supervised Learning for EEG Emotion Recognition. IEEE Trans. Affect. Comput. 14, 2512–2525 (2023).

29. Vecchio, F. et al. Resting state cortical EEG rhythms in Alzheimer’s disease: toward EEG markers for clinical applications: a review. in Supplements to Clinical Neurophysiology vol. 62 223–236 (Elsevier, 2013).

30. Wang, J. et al. Resting state EEG abnormalities in autism spectrum disorders. J. Neurodev. Disord. 5, 24 (2013).

31. Yassine, S. et al. Functional Brain Dysconnectivity in Parkinson’s Disease: A 5-Year Longitudinal Study. Mov. Disord. Off. J. Mov. Disord. Soc. 37, 1444–1453 (2022).

32. De Aguiar Neto, F. S. & Rosa, J. L. G. Depression biomarkers using non-invasive EEG: A review. Neurosci. Biobehav. Rev. 105, 83–93 (2019).

33. Olbrich, S. & Arns, M. EEG biomarkers in major depressive disorder: discriminative power and prediction of treatment response. Int. Rev. Psychiatry Abingdon Engl. 25, 604–618 (2013).

34. Yener, G. et al. Treatment effects on event-related EEG potentials and oscillations in Alzheimer’s disease. Int. J. Psychophysiol. 177, 179–201 (2022).

35. Lenartowicz, A. & Loo, S. K. Use of EEG to Diagnose ADHD. Curr. Psychiatry Rep. 16, 498 (2014).

36. Pavlov, Y. G. et al. #EEGManyLabs: Investigating the replicability of influential EEG experiments. Cortex 144, 213–229 (2021).

37. Trübutschek, D. et al. EEGManyPipelines: A Large-scale, Grassroots Multi-analyst Study of Electroencephalography Analysis Practices in the Wild. J. Cogn. Neurosci. 36, 217–224 (2024).

38. Lefebvre, A. et al. Alpha Waves as a Neuromarker of Autism Spectrum Disorder: The Challenge of Reproducibility and Heterogeneity. Front. Neurosci. 12, 662 (2018).

39. Pascucci, D. et al. EEG brain waves and alpha rhythms: Past, current and future direction. Neurosci. Biobehav. Rev. 176, 106288 (2025).

40. Marek, S. et al. Reproducible brain-wide association studies require thousands of individuals. Nature 603, 654–660 (2022).

41. Thirion, B. On the statistics of brain/behavior associations. Aperture Neuro 83 (2023) doi:10.52294/51f2e656-d4da-457e-851e-139131a68f14.

42. Morales, S. et al. Generalizability of developmental EEG: Demographic reporting, representation, and sample size. Dev. Cogn. Neurosci. 74, 101567 (2025).

43. Westlin, C. et al. Improving the study of brain-behavior relationships by revisiting basic assumptions. Trends Cogn. Sci. 27, 246–257 (2023).

44. Gratton, C., Nelson, S. M. & Gordon, E. M. Brain-behavior correlations: Two paths toward reliability. Neuron 110, 1446–1449 (2022).

45. Ebadi, A. et al. Beyond homogeneity: charting the landscape of heterogeneity in neurodevelopmental and psychiatric electroencephalography. Transl. Psychiatry 15, 223 (2025).

46. Tabbal, J. et al. Characterizing the heterogeneity of neurodegenerative diseases through EEG normative modeling. Npj Park. Dis. 11, 117 (2025).

47. Alexander, L. M. et al. An open resource for transdiagnostic research in pediatric mental health and learning disorders. Sci. Data 4, 170181 (2017).

48. Langer, N. et al. A resource for assessing information processing in the developing brain using EEG and eye tracking. Sci. Data 4, 170040 (2017).

49. McPartland, J. C. et al. The Autism Biomarkers Consortium for Clinical Trials (ABC-CT): Scientific Context, Study Design, and Progress Toward Biomarker Qualification. Front. Integr. Neurosci. 14, 16 (2020).

50. Pelphrey, K. Multimodal Developmental Neurogenetics of Females with ASD. Preprint at 10.15154/4DAT-5683 (2012).

51. Lubba, C. H. et al. catch22: CAnonical Time-series CHaracteristics. Data Min. Knowl. Discov. 33, 1821–1852 (2019).

52. Fortin, J.-P. et al. Harmonization of cortical thickness measurements across scanners and sites. NeuroImage 167, 104–120 (2018).

53. Spearman, C. The proof and measurement of association between two things. Am. J. Psychol. 15, 72–101 (1904).

54. Virtanen, P. et al. SciPy 1.0: fundamental algorithms for scientific computing in Python. Nat. Methods 17, 261–272 (2020).

55. Clarke, A. R., Barry, R. J., McCarthy, R. & Selikowitz, M. Age and sex effects in the EEG: differences in two subtypes of attention-deficit/hyperactivity disorder. Clin. Neurophysiol. 112, 815–826 (2001).

56. Markovska-Simoska, S. & Pop-Jordanova, N. Quantitative EEG in Children and Adults With Attention Deficit Hyperactivity Disorder: Comparison of Absolute and Relative Power Spectra and Theta/Beta Ratio. Clin. EEG Neurosci. 48, 20–32 (2017).

57. Gur, R. C. et al. Age group and sex differences in performance on a computerized neurocognitive battery in children age 8-21. Neuropsychology 26, 251–265 (2012).

58. Eme, R. F. Selective Females Affliction in the Developmental Disorders of Childhood: A Literature Review. J. Clin. Child Psychol. 21, 354–364 (1992).

59. Vallat, R. Pingouin: statistics in Python. J. Open Source Softw. 3, 1026 (2018).

60. Gramfort, A. et al. MEG and EEG data analysis with MNE-Python. Front. Neurosci. 7, (2013).

