## Supplementary Materials for "Robust EEG brain-behavior associations emerge only at large sample sizes"

**Table S1 | Subscores included in our study**

| **Domain** | **ADHD** | **ASD** | **HC** |
|---|---|---|---|
| ADHD Symptoms | SWAN_Inattention_Avg, SWAN_Hyperactivity_Avg, SWAN_Total_Avg, C3SR_Hyperactivity_Raw, C3SR_Inattention_Raw, SDQ_Hyperactivity_Total, CBCL_Attention_Raw | CASI_ADHDInattentive_Severityscore, CASI_ADHDHyperImpulse_Severityscore, CASI_ADHDCombined_Severityscore, CBCL_Attention_Raw | SWAN_Inattention_Avg, SWAN_Hyperactivity_Avg, SWAN_Total_Avg, SDQ_Hyperactivity_Total, CBCL_Attention_Raw, CASI_ADHDInattentive_Severityscore, CASI_ADHDHyperImpulse_Severityscore, CASI_ADHDCombined_Severityscore |
| Autistic Traits | SCQ_Score, ASSQ_Score, GARS_RepBehavior_Raw, GARS_EmotionalResponses_Raw, GARS_CogStyle_Raw, GARS_MaladaptiveSpeech_Raw, GARS_AutismIndex, SRS_Awareness_Raw, SRS_Cognition_Raw, SRS_Motivation_Raw, SRS_RestrictedRepetitive_Raw, SRS_Total_Raw, RBS_Score | ADOS_RestrictedRepetitiveBehavior, ADOS_Total, CASI_Asperger_Severityscore, CASI_Autism_Severityscore, ADI_SectionC_RR_Total, SRS_Awareness_Raw, SRS_Cognition_Raw, SRS_Motivation_Raw, SRS_Total_Raw | SCQ_Score, ASSQ_Score, SRS_Awareness_Raw, SRS_Cognition_Raw, SRS_Motivation_Raw, SRS_RestrictedRepetitive_Raw, SRS_Total_Raw |
| Cognitive Function | WISC_Vocabulary_Raw, WISC_VCI_Raw, WISC_FullScale4_Raw, WISC_BlockDesign_Scaled, WISC_Similarities_Scaled, WISC_Matrix_Scaled, C3SR_LearningProblems_Raw | DAS_RecallDesign_Raw, DAS_WordDefinition_Raw, DAS_PatternConstruction_Raw, DAS_Matrices_Raw, DAS_VerbalSimilarities_Raw, DAS_SequentialQuantitativeReasoning_Raw | WISC_Vocabulary_Raw, WISC_VCI_Raw, WISC_FullScale4_Raw, WISC_BlockDesign_Scaled, WISC_Similarities_Scaled, WISC_Matrix_Scaled |
| Demographics | Age | Age, Barratt_Education, Barratt_Occupation, Barratt_Total | Age, Barratt_Education, Barratt_Occupation, Barratt_Total |

| | | | |
|---|---|---|---|
| Emotional Regulation | DTS_Score, CCSC_ProblemFocusedCoping, CCSC_CogDescisionMaking, CCSC_DirectProblemSolving, CCSC_SeekingUnderstanding, CCSC_AvoidanceCoping, CCSC_AvoidantActions, CCSC_Repression, CCSC_WishfulThinking, CCSC_PosCogRestructing, CCSC_Control, CCSC_Optimism, CCSC_Positive, CCSC_Religion, CCSC_SupportSeeking, CCSC_SupportMotherFather, CCSC_SupportOtherAdults, CCSC_SupportPeers, CCSC_SupportSiblings, PANAS_PositiveAffect | DTS_Score | DTS_Score |
| Executive Function | NIH_final_Card_Sort_Raw, NIH_final_Flanker_Raw, NIH_final_List_Sort_Raw, NIH_final_Processing_Raw, WISC_WorkingMemoryIndex_Raw, NIH_Scores_Card, NIH_Scores_Flanker, NIH_Scores_List, NIH_Scores_Pattern | NIH_final_Card_Sort_Raw, NIH_final_Flanker_Raw, NIH_final_List_Sort_Raw, NIH_final_Processing_Raw, WISC_WorkingMemoryIndex_Raw, NIH_Scores_Card, NIH_Scores_Flanker, NIH_Scores_List, NIH_Scores_Pattern | NIH_final_Card_Sort_Raw, NIH_final_Flanker_Raw, NIH_final_List_Sort_Raw, NIH_final_Processing_Raw, WISC_WorkingMemoryIndex_Raw |
| Externalizing Symptoms | PBS_Score, C3SR_Aggression_Raw, SDQ_Conduct_Problems_Total, SDQ_Difficulties_Total, SDQ_Externalising_Total, CBCL_Rulebreak_Raw, CBCL_Aggressive_Raw, CBCL_External_Raw, CBCL_Total_C_Raw, CBCL_Total_Raw | CASI_OppositionDefiantDis_Severityscore, CASI_ConductDis_Severityscore, CBCL_Rulebreak_Raw, CBCL_Aggressive_Raw, CBCL_External_Raw, CBCL_Total_C_Raw, CBCL_Total_Raw | SDQ_Conduct_Problems_Total, SDQ_Difficulties_Total, SDQ_Externalising_Total, CBCL_Rulebreak_Raw, CBCL_Aggressive_Raw, CBCL_External_Raw, CBCL_Total_C_Raw, CBCL_Total_Raw, CASI_OppositionDefiantDis_Severityscore, CASI_ConductDis_Severityscore |
| Family Environment | APQ_Total, PSI_Score, APQ_P_Total, CPIC_Frequency_Total, CPIC_Intensity_Total, CPIC_Resolution_Total, CPIC_Content_Total, CPIC_Perceived_Threat_Total, CPIC_Self_Blame_Total, CPIC_Triangulation_Total, CPIC_Stability_Total, C3SR_FamilyRelation_Raw | APQ_Total, APQ_P_Total | APQ_Total, APQ_P_Total |

| | | | |
|---|---|---|---|
| Functional Impairment | WHODAS_P_Score, WHODAS_SR_Score, SDQ_Generating_Impact_Total | SDQ_Generating_Impact_Total | SDQ_Generating_Impact_Total |
| General Psychopathology | TRF_Total, YSR_Total | | |
| Global Functioning | CGAS_Score, CIS_SR_Total | | |
| Internalizing Symptoms | MFQ_SR_Score, SCARED_P_Score, SCARED_SR_Score, PANAS_NegativeAffect, SDQ_Emotional_Problems_Total, SDQ_Internalising_Total, CBCL_Anxious_Raw, CBCL_Withdrawn_Raw, CBCL_Somatic_Raw, CBCL_Internal_Raw, MFQ_P_Score | SCARED_P_Score, SDQ_Emotional_Problems_Total, SDQ_Internalising_Total, CBCL_Anxious_Raw, CBCL_Withdrawn_Raw, CBCL_Somatic_Raw, CBCL_Internal_Raw, CASI_GeneralAnxDis_Severityscore, CASI_SeparationAnxDis_Severityscore, CASI_MajorDepressionDis_Severityscore, CASI_DysthymicDis_Severityscore, CASI_SocialPhobia_Severityscore | SCARED_P_Score, SDQ_Emotional_Problems_Total, SDQ_Internalising_Total, CBCL_Anxious_Raw, CBCL_Withdrawn_Raw, CBCL_Somatic_Raw, CBCL_Internal_Raw, CASI_GeneralAnxDis_Severityscore, CASI_SeparationAnxDis_Severityscore, CASI_MajorDepressionDis_Severityscore, CASI_DysthymicDis_Severityscore |
| Language | CTOPP_Elision_Raw, CTOPP_BlendingWords_Raw, CTOPP_NonWordRepetition_Raw, CTOPP_RapidDigitNaming_Raw, CTOPP_RapidLetterNaming_Raw, CTOPP_RapidSymbolicNaming_Sum, GFTA_SoundsInWords_Raw, TOWRE_Score_Raw | ADI_SectionB_NonVerbal_Total, ADI_SectionB_Verbal_Total | CTOPP_Elision_Raw, CTOPP_BlendingWords_Raw, CTOPP_NonWordRepetition_Raw, CTOPP_RapidDigitNaming_Raw, CTOPP_RapidLetterNaming_Raw, CTOPP_RapidSymbolicNaming_Sum, TOWRE_Score_Raw |
| Personality | ARI_SR_Score, ICU_P_Score, GRIT_Score, ICU_SR_Score | ARI_SR_Score, ICU_P_Score, ICU_SR_Score | ARI_SR_Score, ICU_P_Score |
| Psychosis Spectrum | CBCL_Thought_Raw | CASI_Schizophrenia_Severityscore | CBCL_Thought_Raw |
| Response Bias | C3SR_NegativeImpression_Raw, C3SR_PositiveImpression_Raw | | |

| | | | |
|---|---|---|---|
| Social | GARS_SocialInteration_Raw, GARS_SocialCommunication_Raw, SRS_Communication_Raw, SDQ_Peer_Problems_Total, SDQ_Prosocial_Total, CBCL_Social_Raw | ADOS_SocialAffect, ADI_SectionA_Social_Total, SRS_Communication_Raw | SRS_Communication_Raw, SDQ_Peer_Problems_Total, SDQ_Prosocial_Total, CBCL_Social_Raw |
| Stress and Adversity | NLES_P_Upset_Total, NLES_P_Upset_Average, ACE_P_Total, NLES_Upset_Total, NLES_Upset_Average | | NLES_P_Upset_Total, NLES_P_Upset_Average |

**Fig S1 | Participant counts per behavioral score** for a) ADHD, b) ASD, and c) HC

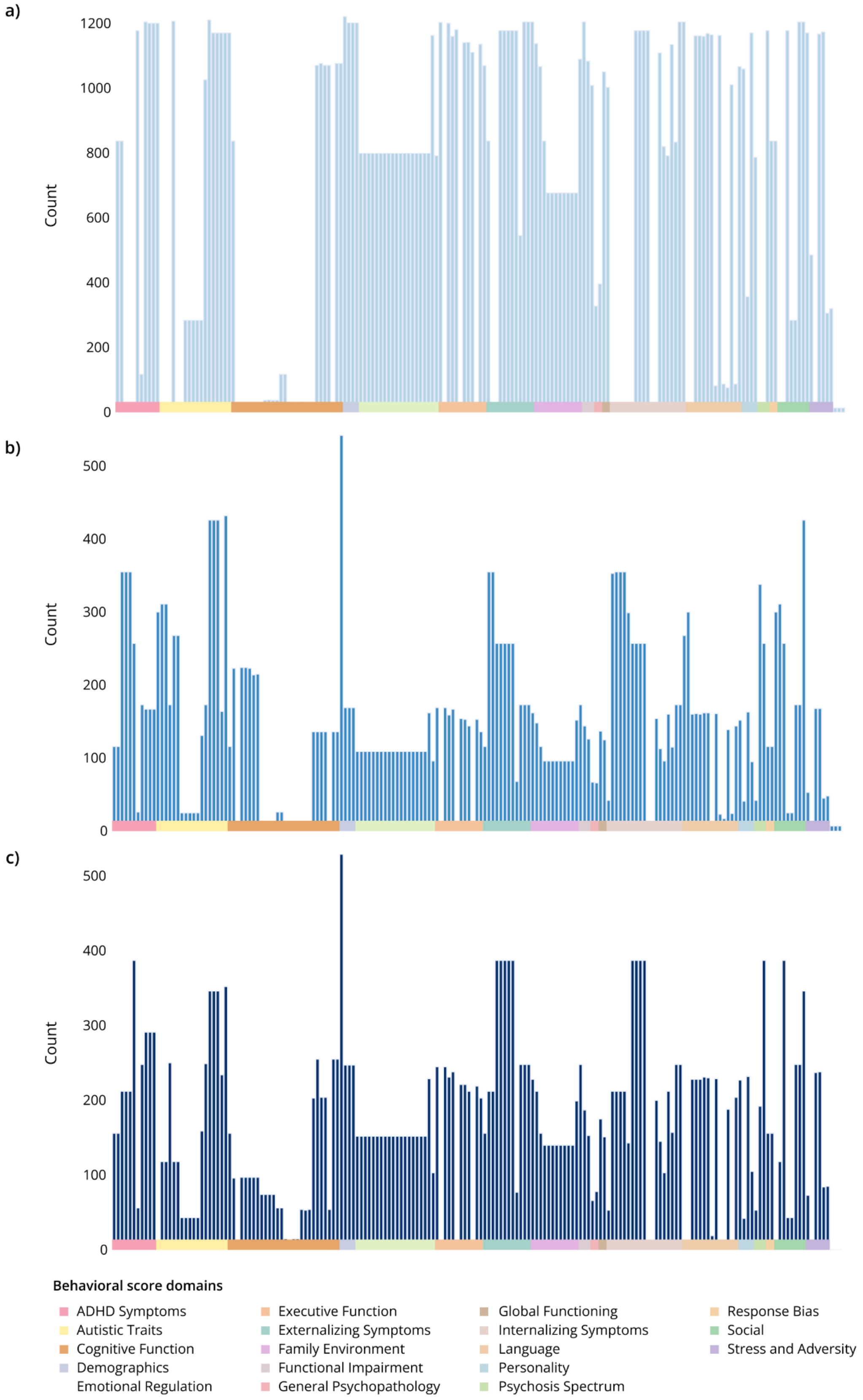

**Table S2 | Summary of features included in this study.** Where relevant, features were computed across multiple frequency bands. For Catch22 features, only the primary category is listed with numbers indicating the count of features from each category. Mean and standard deviation features, which lack specific categories, are not listed here but included in our study.

| Feature Name |
|---|
| **Frequency-domain features** |
| Absolute power (δ, θ, α, β, γ) |
| Relative power (δ, θ, α, β, γ) |
| Alpha peak frequency |
| Theta/beta ratio |
| Theta/alpha ratio |
| Delta/beta ratio |
| Delta/alpha ratio |
| Aperiodic intercept |
| Aperiodic slope |
| Adjusted power (δ, θ, α, β, γ) |
| **Complexity features** |
| Approximate entropy |
| Sample entropy |
| Spectral entropy |
| Hjorth parameters: activity |
| Hjorth parameters: mobility |
| Hjorth parameters: complexity |
| Hurst exponent: H |
| Hurst exponent: c |
| Katz's fractal dimensions (Broadband, δ, θ, α, β, γ) |
| Higuchi's fractal dimension (Broadband, δ, θ, α, β, γ) |
| Detrended fluctuation analysis (Broadband, δ, θ, α, β, γ) |
| Lempel-Ziv complexity (Broadband, δ, θ, α, β, γ) |
| **Time-domain features** |
| Amplitude total power (Broadband, δ, θ, α, β, γ) |
| Mean of the envelope (Broadband, δ, θ, α, β, γ) |
| Standard deviation of the envelope (Broadband, δ, θ, α, β, γ) |
| Skewness of the signal (Broadband, δ, θ, α, β, γ) |
| Kurtosis of the signal (Broadband, δ, θ, α, β, γ) |
| **Catch22 features** |

| |
|---|
| Distribution shape (2) |
| Extreme event timing (2) |
| Incremental differences (2) |
| Linear autocorrelation (4) |
| Linear autocorrelation structure (2) |
| Nonlinear autocorrelation (2) |
| Other |
| Self-affine scaling (2) |
| Simple forecasting |
| Symbolic (4) |

**Table S3 | Subsample size scale for each group.** Scales were generated manually according to the number of available subjects per group

| Group | Subsample sizes (n) |
|---|---|
| **HC/ASD** | 10, 20, 30, 40, 50, 70, 90, 120, 150, 180, 210, 240, 270, 300, 350, 400, 450, 500, 550 |
| **ADHD** | 10, 20, 30, 40, 50, 70, 90, 120, 150, 180, 210, 240, 270, 300, 350, 400, 450, 500, 550, 600, 650, 700, 750, 800, 850, 900, 950, 1000, 1050, 1100, 1200 |

**Fig S2 | Heatmap of thresholds across all associations** for a) ADHD, b) ASD, c) HC

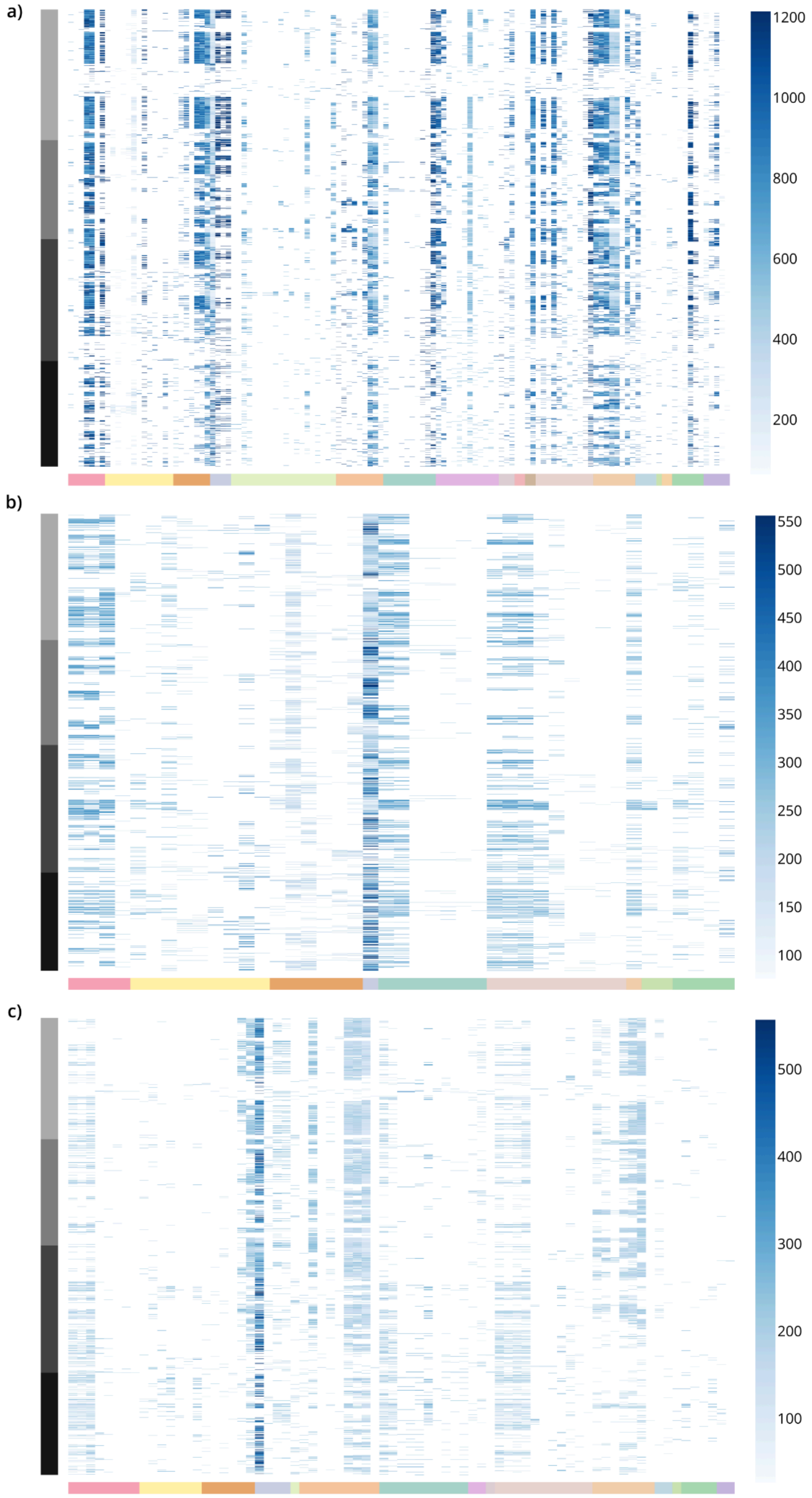

**Fig. S3 | Sample size thresholds across channels.** Sample size thresholds varied across channels for the same feature–score association. As an example, all associations with the CBCL Withdrawn score are shown.

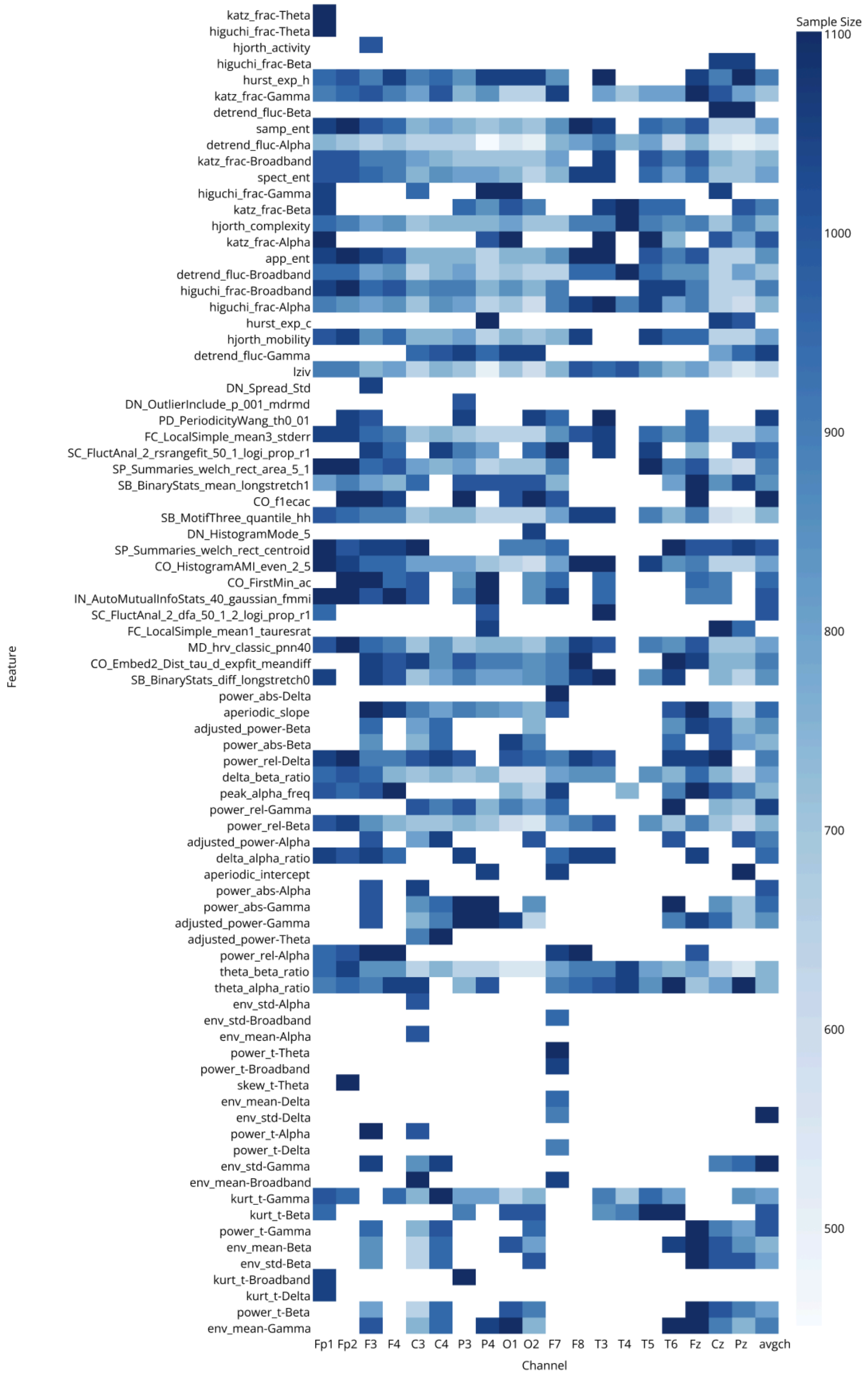

**Fig S4. Error properties and effect size inflation.** For a) ADHD (without limiting the number of participants per score to 1000), b) ASD, c) HC

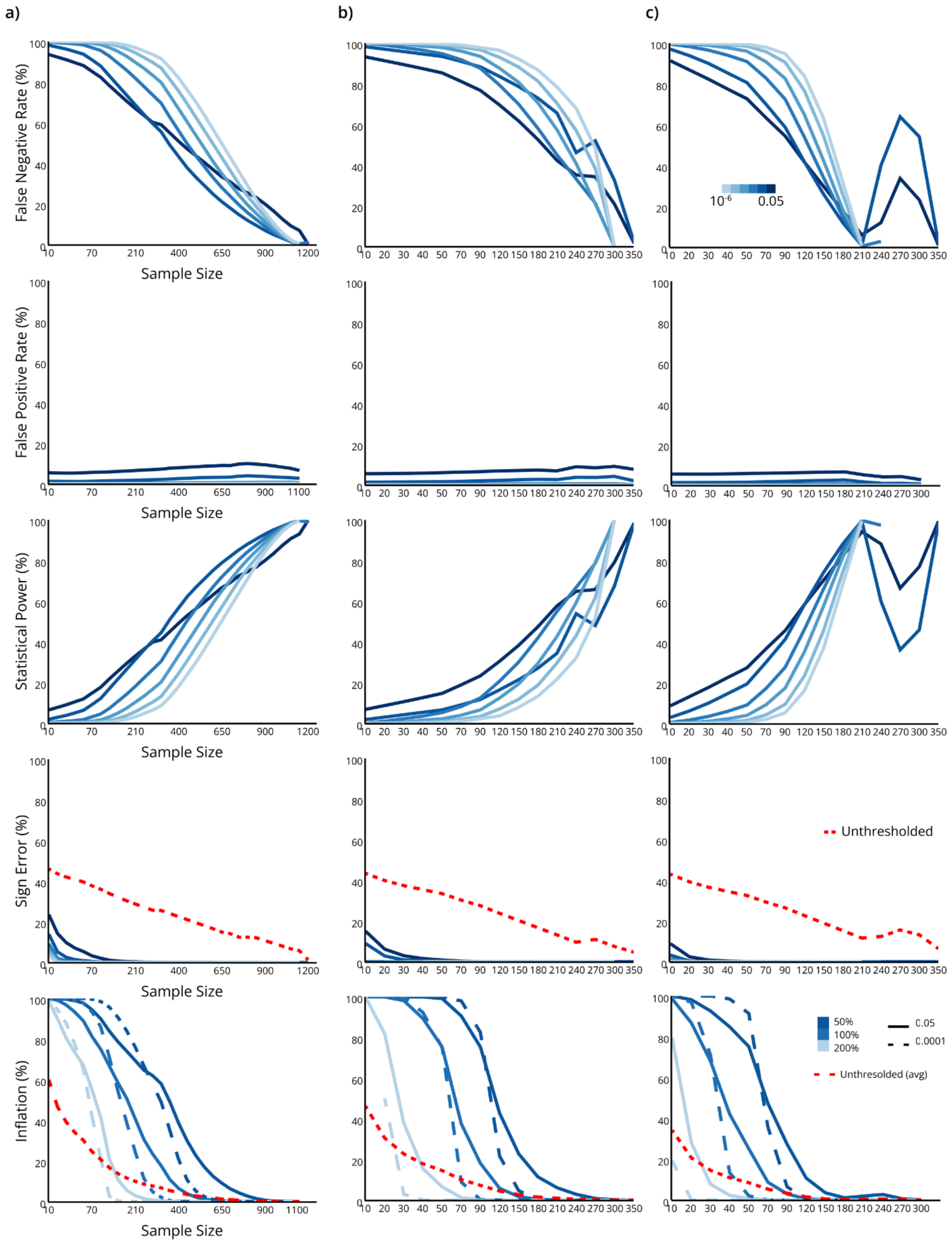

**Fig S5. Error properties and effect size inflation for Spearman partial correlation.** For ADHD (without limiting the number of participants per score to 1000)

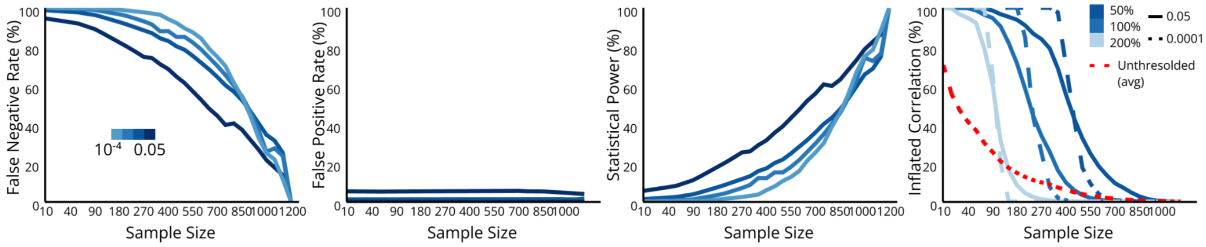

**Fig. S6. Spearman correlation vs Partial correlation.** The figure demonstrates effect sizes at full sample size (left) and distribution of sample size thresholds (right) for a) ASD, b) HC

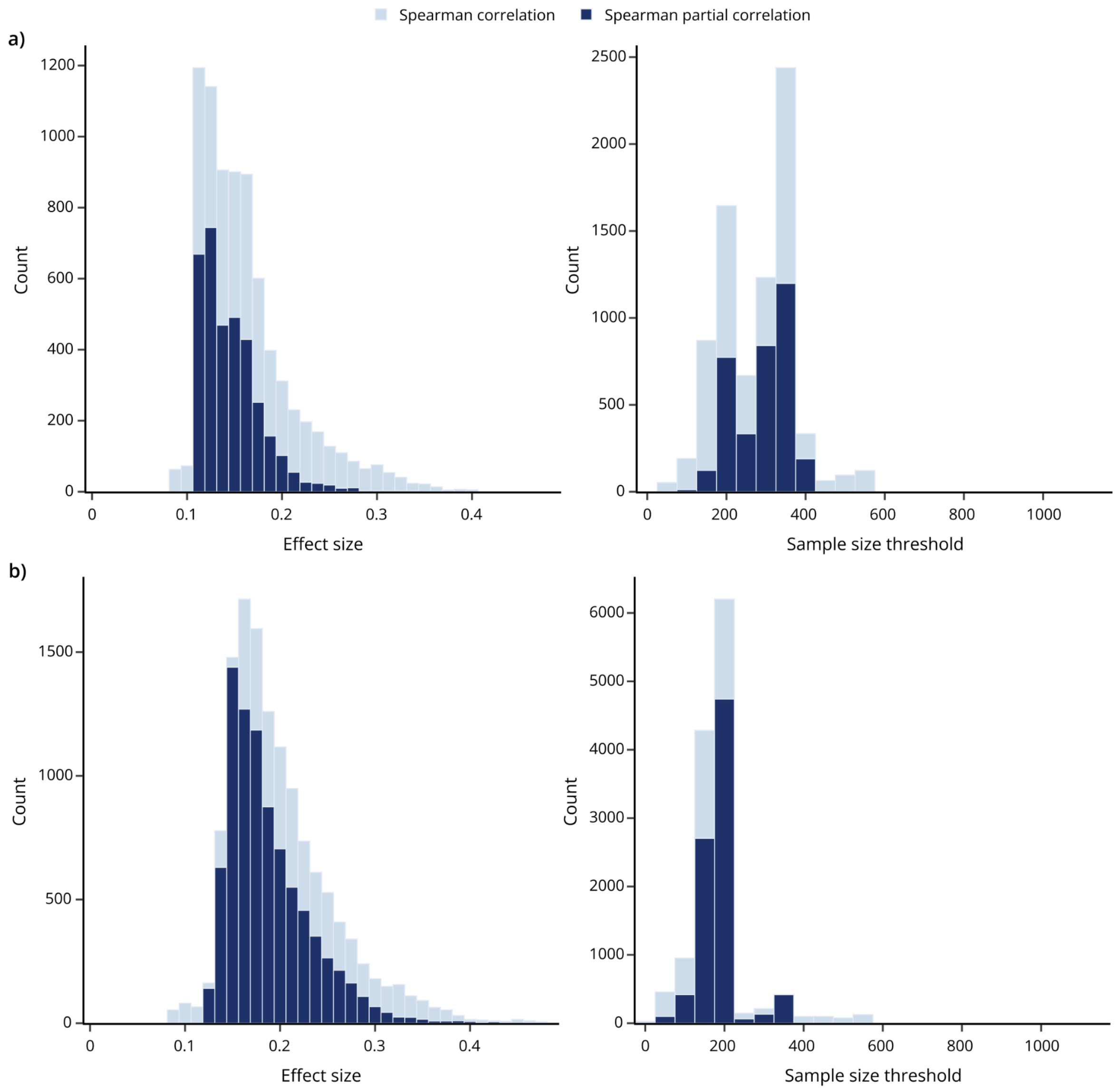
